# Improving protein function prediction by learning and integrating representations of protein sequences and function labels

**DOI:** 10.1101/2024.03.11.584495

**Authors:** Frimpong Boadu, Jianlin Cheng

## Abstract

**Motivation:** As fewer than 1% of proteins have protein function information determined experimentally, computationally predicting the function of proteins is critical for obtaining functional information for most proteins and has been a major challenge in protein bioinformatics. Despite the significant progress made in protein function prediction by the community in the last decade, the general accuracy of protein function prediction is still not high, particularly for rare function terms associated with few proteins in the protein function annotation database such as the UniProt.

**Results:** We introduce TransFew, a new transformer model, to learn the representations of both protein sequences and function labels (Gene Ontology (GO) terms) to predict the function of proteins. TransFew leverages a large pre-trained protein language model (ESM2-t48) to learn function-relevant representations of proteins from raw protein sequences and uses a biological natural language model (BioBert) and a graph convolutional neural network-based autoencoder to generate semantic representations of GO terms from their textual definition and hierarchical relationships, which are combined together to predict protein function via the cross-attention. Integrating the protein sequence and label representations not only enhances overall function prediction accuracy over the existing methods, but substantially improves the accuracy of predicting rare function terms with limited annotations by facilitating annotation transfer between GO terms.

**Availability:** https://github.com/BioinfoMachineLearning/TransFew

**Supplementary information:** Supplementary data are available .

## 1. Introduction

Proteins are essential molecules that play critical functional roles in biological systems. Their functions encompass catalyzing biochemical reactions, serving as structural elements, transducing cellular signals, defending against viruses, regulating gene activities, among others. Elucidating protein functions is crucial for gaining valuable insights into the molecular intricacies of biological systems. However, experimentally determining protein function is a time consuming and laborious process. Currently, fewer than 1% known proteins have function information determined experimentally according to the statistics in UniProt[1]. Therefore, it is important to develop computational methods to predict protein function from sequence and other relevant information.

In the realm of protein function prediction, there are two common challenges: (1) effectively integrating diverse information sources, such as protein sequence, protein-protein interaction, structural features, domain features, and biological texts, to accurately predict protein functions [2], and (2) accurately assigning rare or novel Gene Ontology(GO) terms (labels) [3, 4] with few/no observations in labeled protein function datasets to new proteins that may have the function. It is harder to predict rare (low-frequency) GO terms than common GO terms because the former is less represented than the latter in the function datasets. But it is important to predict rare GO terms because they are usually specific and highly informative function classes that are more useful for generating biological hypotheses than common ones. Moreover, a large portion of all the GO terms are rather rare. Out of over 40, 000 GO terms in the three main Gene Ontology categories: Cellular Component (CC), Molecular Function (MF), and Biological Process (BP), around 20, 000 terms each are assigned to fewer than 100 proteins experimentally [5]. Therefore, there is an urgent need to develop computational methods to predict rare function terms for proteins whose function is described by them.

Predicting rare GO terms is analogous to the few-shot learning problems [6] in various domains like computer vision[7, 8, 9], and natural language processing(NLP). For example, in the classification task of named entity typing[10, 11] in NLP, assigning rare entity types to entity names pose a similar challenge, due to the increasing size and granularity of entity types. Two kinds of methods, i.e., embedding-based methods and generative methods, have been proposed to tackle this challenge[12]. Embedding-based methods focus on learning an embedding space associating low-level features of highly annotated classes with semantic information of both highly annotated classes and rarely annotated classes to transfer knowledge from highly annotated classes to rarely annotated ones with few annotations. Generative methods generate features for rare classes based on samples from adequately annotated classes, converting the problem into the conventional supervised learning. In the protein function prediction, the hierarchical structure and textual descriptions of GO terms (classes/labels) provides us with the vital semantic information to transfer knowledge from the well-annotated classes to the ones with few or no annotations [13].

In this study, we introduce an embedding-based deep learning method called TransFew to predict protein functions[2], with an emphasis on improving the prediction of protein function described by rare GO terms. TransFew generates a function-relevant representations of a single protein sequence in the sequence space using a pretrained protein language model (i.e., ESM2[14, 2]) and multi-layer perceptrons (MLP). The sequence representation of a protein is generated by multiple MLP modules with residual connections each designed to predict functions for proteins in terms of a specific group of GO terms with similar annotation frequency, which therefore cover all the GO terms from rare ones to common ones equally. TransFew also generates a semantic representation of all the GO terms (labels) in the label space from their textual description (definition) and their hierarchical relationships in the Gene Ontology graphs (e.g., the inheritance and composition relationships (i.e., similarity) between GO terms) using a graph convolutional neural network (GCN)-based auto-encoder and a biological natural language model (BioBert) [15, 16], which facilitates the transfer of annotations from common GO terms to rare ones according to their relationships. TransFew then uses a joint feature label embedding technique based on the cross attention to integrte the label representations and sequence representations to accurately predict protein functions.

TransFew not only improves the overall accuracy of protein function prediction over several existing sequence-based function prediction methods, but also substantially enhances the prediction of rare GO terms.

## 2. Materials and Methods

The overall architecture of TransFew is illustrated in Figure 1. It has three components: (1) a query processor consisting of multiple MLPs to extract function-relevant sequence representations from a protein sequence (query), (2) a label processor to extract label representations for all the GO terms (labels), and (3) a joint feature-label embedding network to combine sequence and label representations to predict the function of a protein. One TransFew model was trained to predict the GO terms in each of the three GO function categories (molecular function (MF), cellular component (CC) and biological process (BP)), respectively.

**Fig. 1.**
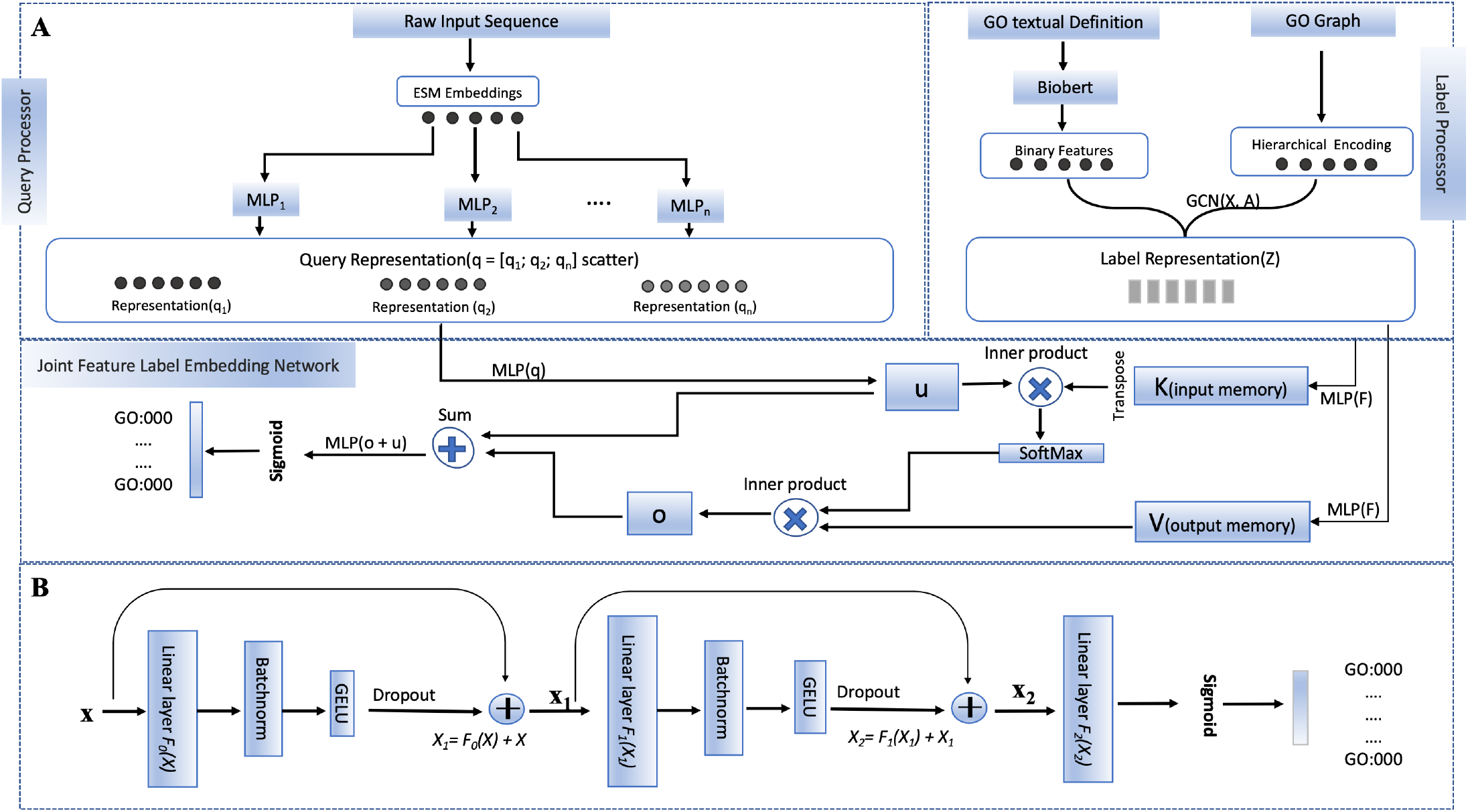
The overall architecture of TransFew. (A) The three components of TransFew: a query processor(top left) to generate sequence representation using multi-layer perceptrons (MLPs) and ESM2, a label processor(top right) to extract label representations using Biobert [17] and a graph convolutional neural networks (GCN)-based auto-encoder[18], and a joint feature-label embedding network to combine sequence and label representations via the cross attention for a MLP to make final function prediction. (B) The detailed design of a typical MLP module used in TransFew.

### 2.1. Query processor

The query processor is to generate the function-relevant sequence representations for proteins. Protein function terms have very different annotation frequency in the labeled protein function datasets. Here, the annotation frequency of a GO term is the number of proteins that are labelled to have it as function. Rare GO terms are the ones that only occur to be the function labels of a small number of proteins. Generating a simple representation for all the GO terms together regardless of their frequency allows the commons GO terms dominate the rare (low-frequency) GO terms, which can reduce the accuracy of predicting them. Therefore, we partitioned GO terms into *n* groups for a Gene Ontology category (i.e., MF, CC or BP) based on their annotation frequency, and design *n* MLPs to learn a representation for the *n* group separately (Figure 1A), which allows each MLP to focus on learning a representation in terms of the GO terms in each group including the ones consisting of rare GO terms. Specifically, the GO terms of BP were partitioned into three groups and the GO terms of CC and MF into two groups. The statistics for the partitions is shown in Table 1

**Table 1.** The partition of GO terms into groups according to their annotation frequency for three gene ontologies (BP, MF, and CC). MF and CC terms are partitioned into two groups according to the frequency threshold of 30 respectively, while BP terms are partitioned into three groups because BP has many more GO terms than MF and CC. The last column reports the number of GO terms in each partition.

| Ontology | Partition | Number of GO Terms |
| --- | --- | --- |
| BP | 1 - 5 | 7892 |
| BP | 6 - 30 | 6977 |
| BP | > 30 | 6415 |
| MF | 1 - 30 | 6040 |
| MF | > 30 | 1183 |
| CC | 1 - 30 | 2083 |
| CC | > 30 | 873 |

Each MLP (i.e., MLP_*i*_) takes as input the sequence features of a protein generated by a large pretrained protein language model, ESM2_t48[14] from its sequence, and outputs a vector 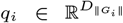, where ∥*G*_*i*_∥ is the number of GO terms in a GO group *G*_*i*_. ESM-2t48 [14] accepts the sequence of a protein as input and generates feature embeddings at multiple layers. Here, the per-residue embeddings of the last layer (48th layers) are taken out and averaged by the mean aggregator to generate the embedding of the protein, whose dimension is 5120. For a protein sequence exceeding the length limit of ESM2_t48, i.e., 1022 residues, it is divided into chunks of length 1022 except the last chuck that may have fewer than 1022 residues, each of which is processed by ESM2_t48 separately. The embeddings for all the chunks are concatenated as the embedding of the full protein sequence. In addition to using ESM-2t48 to generate input features for the MLP, we also tried to use multiple sequence alignments (MSAs) [19] and InterPro domain annotations [20, 21, 22, 23] of proteins to generate input features for the MLPs. The details of generating MSAs and InterPro domain annotations are described in Supplementary Note 1 and Supplementary Note 2. However, according to the ablation study, adding them on top of the features based on ESM-2t48 does not improve protein function prediction accuracy, and therefore they are not included into the final version of TransFew.

The detailed architecture of a MLP of generating the representation of a protein from its sequence features is depicted in Figure 1B. The MLP has multiple blocks, each of which has a fully connected linear layer, followed by a batch normalization layer and a Gaussian Error Linear Unit (GELU). The input for each block except for the last one is added to its output via a skip connection, resulting in a residual network. The output of the last block is used as input for a sigmoid function to predict the probability of each GO term represented as logit.

The entire query processor, along with all other components, is jointly trained. Each MLP is specifically tailored to predict the GO terms within its corresponding protein group prior to integration with other TransFew components. The output *q*_*i*_ (a vector of predicted logits of the GO terms in a group *G*_*i*_) that a MLP generated for an input protein is the representation of the protein in terms of the GO terms in the group. The representations from all the MLPs are combined as the final query representation of the protein in terms all the GO terms in a gene ontology (MF, CC, or BP), which is the output of the query processor.

The combination process involves employing a scatter operation[24, 25], wherein the values produced by each MLP are distributed within the query representation tensor to match the predefined order of the GO terms.

### 2.2. Label processor

The label processor in Figure 1A is used to generate semantic representations for all the GO terms (labels) under consideration. Two types of label data, i.e., the relations between GO terms in a GO Graph and the definition of GO terms (the textual descriptions) are used as input for the label processor.

The relationships between GO terms (nodes) in a GO graph are represented by an adjacency matrix *A*, where each row encodes the relationships of a node. The entry *A*_*ij*_ is set to 1 if node *i* is an ancestor of *j* or equal to *j*, and 0 otherwise. *A* encodes the hierarchical relationships between the GO terms.

For the definitions of the GO terms, we collected the textual description of each GO term, which contains what the term represents as well as reference(s) to the original source of the information. The textual description of each GO term is used by a pre-trained biomedical language model, BioBert [15, 16], to generate an embedding for it. The dimension of the embedding (*D*_*e*_) is 768, which is set by BioBERT. The embedding is considered the semantic features of each GO term.

The hierarchical relationships and the semantic embeddings of the GO terms are integrated by a graph auto-encoder model [18] to generate the representation of all the GO terms (labels). The input for the model is a GO graph, in which the relationships between nodes (GO terms) are stored in the matrix *A* and the feature of each node is its semantic embedding generated from the textual description of its GO term. The model uses an encoder-decoder architecture, where the encoder is a two-layer graph convolutional network (GCN)[26] defined as:

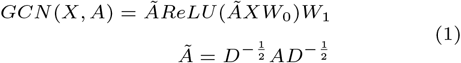

*W*_1_ and *W*_2_ are the weight matrices, Ã is the symmetrically normalized form of the *A*, and *X* is the matrix of the semantic embeddings of all the GO terms. *ReLU* denotes the ReLU activation function. We use the inner product decoder to reconstruct *A* as Â from the embeddings *Z* outputted by the GCN model as follows:

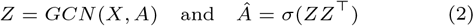

where *σ*(*·*) is the logistic sigmoid function. The graph auto-encoder model was pretrained to reconstruct the GO Graph, *A*, from *A* itself and the semantic embeddings of the GO terms, through the self-supervised learning. After the training, the 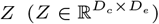 extracted from the bottleneck layer of the GCN-based autoencoder is used as the label representation, where *D*_*c*_ is the number of GO terms and *D*_*e*_ represents the dimension of the label representation (in this work *D*_*e*_ = 1024). It is worth noting that the label representation is independent of any protein.

Additionally, we investigated two alternative encoder architectures, such as Graph Attention Networks (GAT) [27, 28] and Graph Transformer (TransformerConv) [29] to combine the features of the textual description and GO term relationships, but they did not perform better than the the GCN-based auto-endcoder (see Supplementary Note 4).

### 2.3. Joint feature-label embedding network

We designed a joint-label embedding network to match and combine the sequence representation of a protein generated by the query processor and the label representation of all the GO terms generated by the label processor to predict the GO terms for the protein (Figure 1A).

Different from one previous work [30] using a bilinear function to score the match between a given protein query (*q*) and ontology terms and another work [13] applying a scoring function based on softmax and 1D convolutional network to match them, we developed a cross attention-based joint embedding model to measure the association between query proteins and GO terms to improve protein function prediction.

Given the label representation 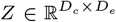, and the query protein representation 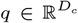, TransFew converts the query representation *q* to 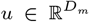 using a linear layer as follows: 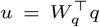, and 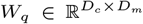, and constructs two memory components: key 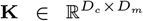 and value 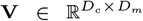 from Z, using two embedding matrices 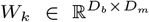 and 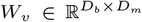 respectively. The cross attention between the representation of a query protein *q* and the representation of all the GO term (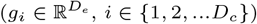, *i ∈ {*1, 2, …*D*_*c*_*}*) is computed as:

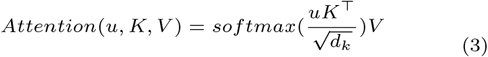

where *d*_*k*_ = *D*_*m*_.

The representation of the query protein and the cross attention are combined by a MLP with a residual connection to predict the probability of GO terms (*y*) for the query protein as follows:

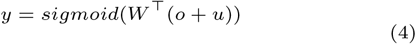

where 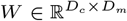 and *o* = *Attention*(*u, K, V*)

The entire model (Figure 1A) including the feature-label embedding network, the query processor, and the pretrained label processor was optimized by minimizing the binary cross-entropy loss between predictions and true labels. It is worth noting that the protein function prediction problem is a multi-label classification problem, in which a protein may have multiple correct labels.

### 2.4. Data Sets

We collected proteins from the UniProtKB/Swiss-Prot data repository that were released by November 2022 for training and validation. The proteins were split into the training dataset and test dataset according to the 90% - 10% ratio. The functional annotations (GO terms) of the proteins were obtained from from UniProt, and the GO ontology graph as well as GO textual data were collected from the Gene Ontology Resource[31, 4]. To get all the terms (labels) associated with a protein, we first retrieved its immediate GO terms provided in UniProt and then for each immediate GO term we traveled up the GO graph to retrieve all its ancestor GO terms. Only the GO terms with relatively strong evidence codes: EXP, IDA, IPI, IMP, IGI, IEP, TAS, IC, HTP, HDA, HMP, HGI, HEP are used as the function labels for each protein, following the criteria used in the Critical Assessment of Functional Annotation (CAFA) [32].

To create an independent test dataset, we obtained proteins in the UniProtKB/Swiss-Prot database whose function annotation were released in December 2023. This test dataset is called Test_all. Moreover, we used MMseqs [33] to filter out the sequences in Test_all that have more than 30% identity with the sequences in the training dataset to create a redundancy reduced dataset - Test_novel, which is used to test how well TransFew can generalize to new proteins that have little or no sequence similarity with the training proteins.

The number of proteins in the training dataset, validation dataset, Test_all dataset, and Test_novel dataset for each gene ontology category (MF, CC, and BP) is reported in Table 2.

**Table 2.**
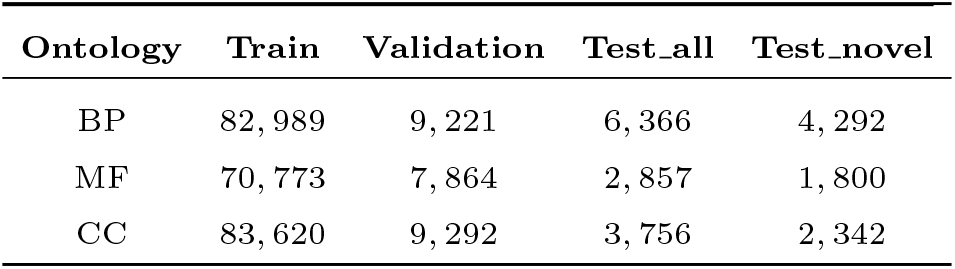
The number of proteins in training, validation, Test_all, and Test_novel datasets for each of the three GO categories (BP, MF, and CC).

| Ontology | Train | Validation | Test_all | Test_novel |
| --- | --- | --- | --- | --- |
| BP | 82,989 | 9,221 | 6,366 | 4,292 |
| MF | 70,773 | 7,864 | 2,857 | 1,800 |
| CC | 83,620 | 9,292 | 3,756 | 2,342 |

## 3. Results & Discussions

### 3.1. Benchmarking TransFew with three baseline methods on the test datasets

We compared TransFew with three baseline methods (Naïve, Tale, and NetGO3[34, 22, 21]) on the Test_all dataset in terms of multiple metrics of evaluating protein function prediction, including F_*max*_, Area under the Precision-Recall curve (AUPR), weighted F_*max*_, and S_*min*_ of measuring the uncertain/missing information in function predicitons [35, 36, 37] (see the detailed definition of the evaluation metrics in Supplementary Note 5).

The Naïve method simply uses the frequency of Gene Ontology (GO) terms in the training dataset to make predictions.

Tale [38] is a transformer-based method that integrates protein sequence and label features to predict protein function by jointly embedding sequence and hierarchical label information. Although Tale and TransFew are common in combining sequence and label features, they have several significant differences. Firstly, TransFew incorporates both hierarchical relationships and textual definitions to generate label representations, while Tale only uses the former. Secondly, TransFew extracts sequence embeddings of a protein using a pre-trained language model (ESM2), whereas Tale uses its own encoder to generate the embeddings. Thirdly, the MLPs to generate sequence representations of proteins in terms of both common and rare GO terms, the GCN-based auto-encoder of generating label representations, and the cross attention mechanism of combining the sequence and label representations in TransFew are all different from the network architecture used in Tale. It is also worth noting that Tale combines its machine learning predictions with the homology-based function predictions made by DIAMOND [39] to further improve its performance. We ran a locally installed Tale to obtain its predictions for the test proteins.

NetGO3[34, 22, 21] is a highly sophisticated ensemble method combining the outputs of seven individual function prediction methods using different sources of input information: Naive prediction based on GO term frequency, BLAST-KNN based on k-nearest neighbors of BLAST search), LR-3mer based on the logistic regression of the frequency of amino acid trigrams, LR-InterPro based on the logistic regression of InterPro features encompassing rich domain, family, and motif information, Net-KNN which extracts and incoporates protein-protein interaction network information from STRING database[40] into the system, LR-Text utilizing the text data about proteins extracted from PubMed[41], and LR-ESM that generates embeddings for each protein using ESM-1b[42] for a learning-to-rank algorithm to predict function. We utilized the NetGO3 web server to obtain its predictions for the test proteins.

The results of TransFew, Naive, Tale and NetGO3 on the Test_all dataset are presented in Table 3A. TransFew performs best in all three GO categories in terms of all the metrics among all the situations except one. In some cases, the improvement over the second best performing method is substantially. For instance, Fmax of TransFew for predicting GO terms in BP is 0.605, 40% higher than 0.431 of NetGO3. The precision-recall curves of the four methods for the three gene ontology categories (BP, MF, and CC) on the Test_all dataset are visualized in Figure 2 respectively. The outermost curve above all the other curves is that of TransFew. Its improvement of predicting the GO terms in BP is particularly substantial.

**Table 3.** The performance of TransFew, Naive, Tale, and NetGO3 on the test datasets in the three GO categories (BP, MF, and CC). (A) The results on all the new proteins in Test_all. (B) The results on Test_novel comprised of proteins that have *≤* 30% sequence identity with the proteins in the training dataset of TransFew. Bold font highlights the best result. TransFew was trained using all the GO terms with at least one annotation in the training dataset.

| (A) On Test_all |  |  |  |  |  |  |  |  |  |  |  |  |
| --- | --- | --- | --- | --- | --- | --- | --- | --- | --- | --- | --- | --- |
| Methods | $F_{max}$ (↑) | | | $WF_{max}$ (↑) | | | $AUPR$ (↑) | | | $S_{min}$ (↓) | | |
|  | BP | MF | CC | BP | MF | CC | BP | MF | CC | BP | MF | CC |
| Naive | 0.246 | 0.374 | 0.539 | 0.209 | 0.309 | 0.349 | 0.127 | 0.143 | 0.425 | 32.826 | 10.986 | 9.779 |
| Tale | 0.290 | 0.581 | 0.648 | 0.252 | 0.554 | 0.529 | 0.169 | 0.530 | 0.688 | 32.786 | 8.109 | 8.104 |
| NetGO3 | 0.431 | 0.659 | 0.696 | 0.397 | 0.637 | 0.597 | 0.352 | **0.616** | 0.669 | 28.278 | 6.749 | 7.146 |
| TransFew | **0.605** | **0.717** | **0.736** | **0.585** | **0.703** | **0.643** | **0.501** | 0.555 | **0.734** | **18.024** | **5.483** | **6.379** |
| (B) On Test_novel |  |  |  |  |  |  |  |  |  |  |  |  |
| Methods | $F_{max}$ (↑) | | | $WF_{max}$ (↑) | | | $AUPR$ (↑) | | | $S_{min}$ (↓) | | |
|  | BP | MF | CC | BP | MF | CC | BP | MF | CC | BP | MF | CC |
| Naive | 0.245 | 0.367 | 0.552 | 0.206 | 0.321 | 0.365 | 0.121 | 0.137 | 0.434 | 31.439 | 10.366 | 9.312 |
| Tale | 0.289 | 0.563 | 0.659 | 0.252 | 0.540 | 0.539 | 0.168 | 0.496 | 0.703 | 31.296 | 7.812 | 7.719 |
| NetGO3 | 0.423 | 0.632 | 0.707 | 0.391 | 0.616 | 0.602 | 0.342 | **0.565** | 0.674 | 27.150 | 6.684 | 6.756 |
| TransFew | **0.567** | **0.667** | **0.732** | **0.543** | **0.653** | **0.634** | **0.461** | 0.494 | **0.719** | **18.638** | **6.061** | **6.338** |

**Fig. 2.**
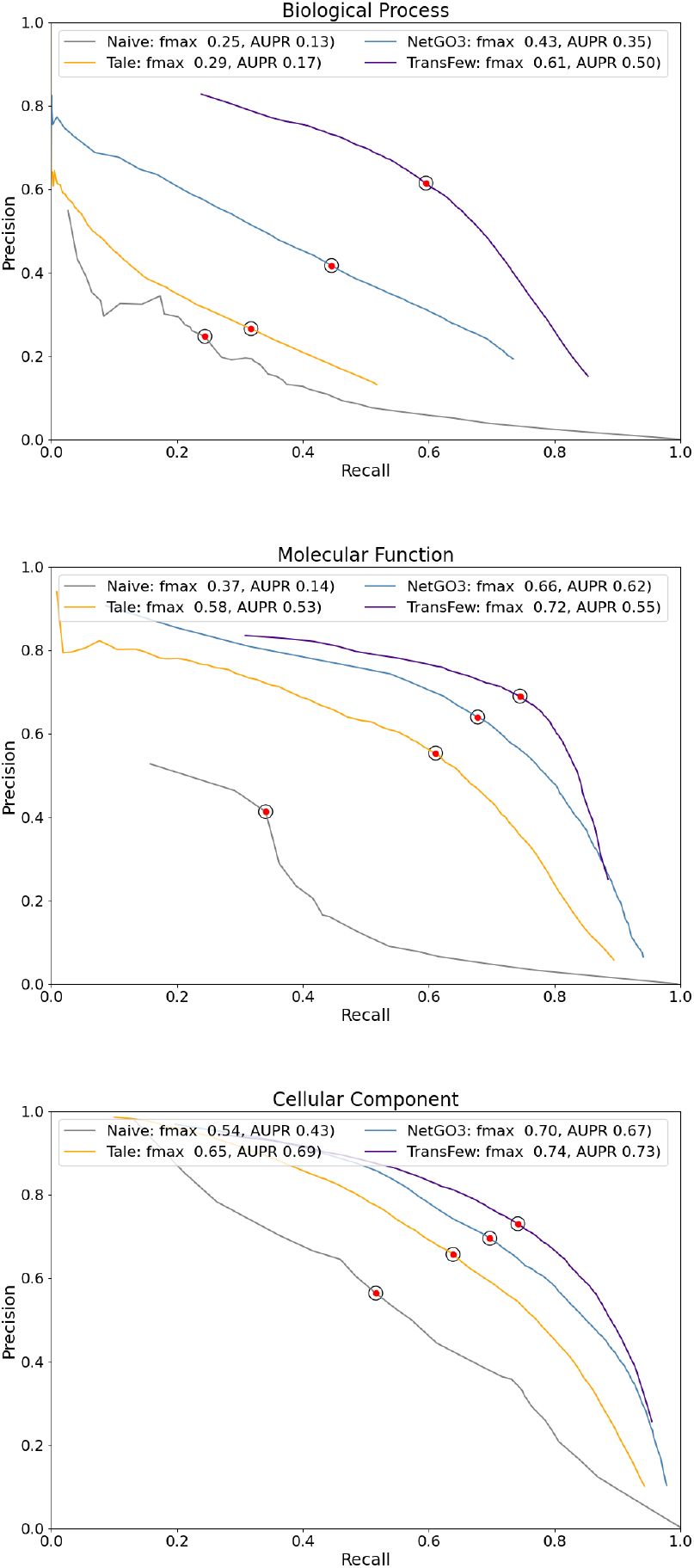
The Precision-Recall curves of TransFew, Naive, Tale and NetGO3 for the three ontologies (BP, MF, and CC) on the Test_all dataset, respectively. The circled red dot highlights the point where each method achieves the highest *F*_*max*_

On the Test_novel dataset consisting of proteins that have *≤* 30% sequence identity with the proteins in the training data, TransFew also performs best among all the methods in all three GO categories among all the situations except one (Table 3B). The performance of TransFew on Test_novel is only slightly or moderately lower than on Test_all in terms of different metrics, indicating that it generalizes well to new test proteins that have no or little sequence identity with the training proteins.

It is worth noting that TransFew is a pure machine learning method, while NetGO3 and Tale combines machine learning predictions and homology-based function annotation transfer to make final prediction. The results show that a pure end-to- end machine learning method like TransFew can perform better than ensemble methods based on both machine learning and homology transfer for protein function prediction.

### 3.2. Performance of predicting rare GO terms

We investigated how well TransFew predicted the rare GO terms with low frequency (few annotations). We group GO terms with *<*= 100 annotations into 20 groups according to their number of annotations (frequency) in the training data at an interval size of 5. The average AUPRC scores of TransFew, Tale, and NetGO3 predicting the GO terms in each group for BP, MF, and CC are shown in Figure 3. TransFew outperforms Tale for all the groups and NetGO3 for most groups. Particularly, its improvement of predicting GO terms in BP over Tale and NetGO3 is very substantial. The average AUPR score of TransFew predicting rare GO terms in BP is usually a few times that of Tale and NetGO3. For the rarest GO terms with *≤* 5 annotations, the AUPR score of TransFew is also substantially higher than NetGO3 and Tale for BP, MF, and CC.

**Fig. 3.**
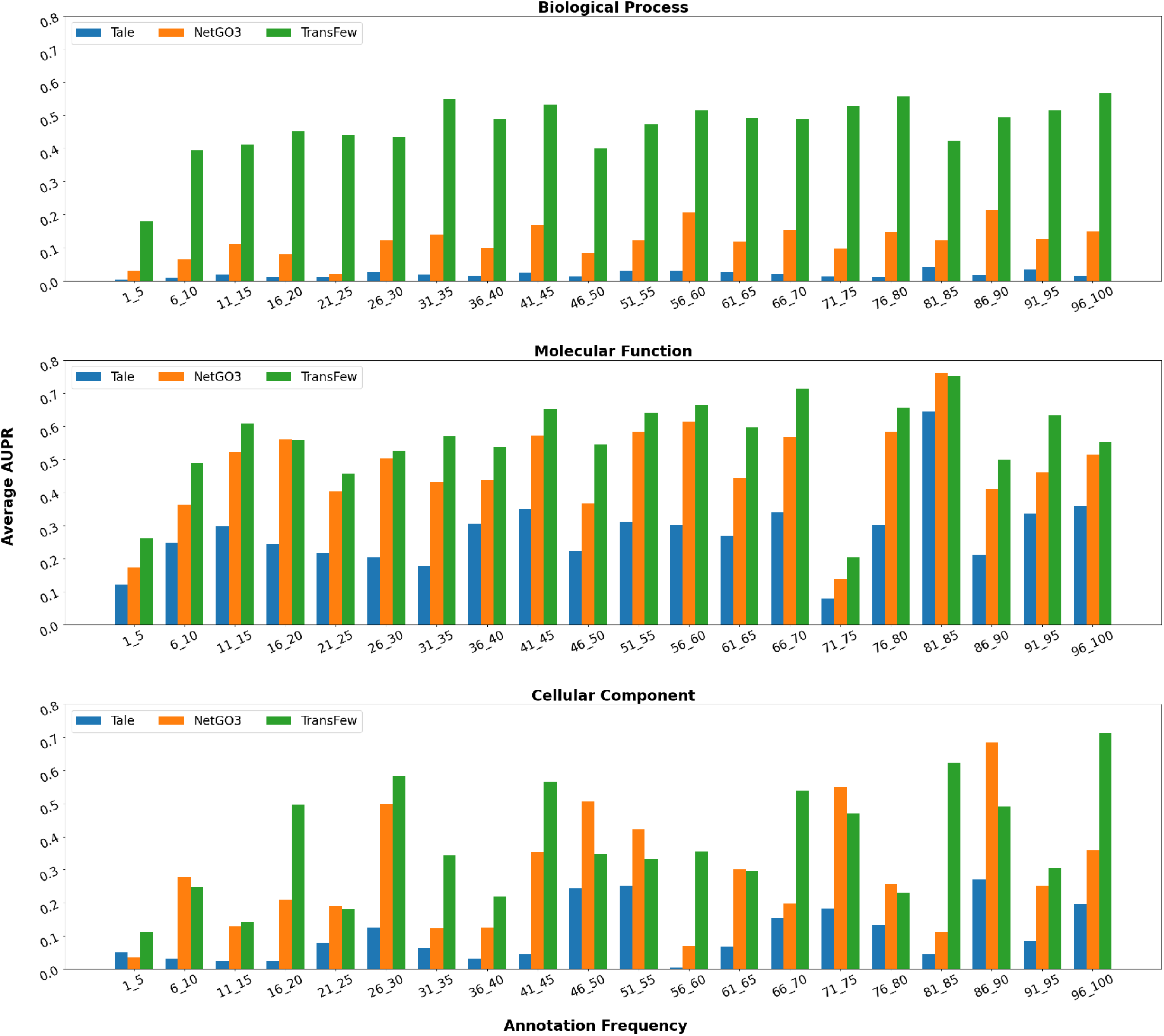
The AUPR score of three methods (TransFew, Tale, and NetGO3) predicting relatively rare GO terms in each GO group within an interval of annotation frequency for the three gene ontologies (BP, MF, and CC). The X-axis represents the annotation frequency of the GO term groups. The Y-axis represents the average AUPR.

The AUPR score of TransFew for different GO term groups only moderately oscillates with respect to their annotation frequency. The Pearson’s correlation between the AUPR of TransFew and the annotation frequency of the GO terms in BP, MF, CC, MF and BP is only 0.20, 0.33, and 0.28, respectively. The results indicate its performance is rather robust with the respect to the change of the annotation frequency of GO terms.

### 3.3. The contributions of different components and implementations of TransFew

We tested how different components or implementations of TransFew influenced its performance. MLP (Interpro), MLP (MSA), and MLP (ESM) denote the three implementations of using the Interpro domain features, the MSA features, and the sequence features generated by ESM2_t48 respectively to generate the sequence representation for function prediction, without using the label representation at all. TransFew stands for the final implementation that combines the sequence representation generated from the ESM2_t48 features and the label representation to predict protein function. TransFew + MSA + Interpro is the same as TransFew except that it uses ESM2_t48 features together with the MSA and Interpro features to generate the sequence representation. MLP (Interpro), MLP (MSA), and MLP (ESM) were trained on the GO terms that have at least 30 annotations, while TransFew and TransFew + MSA + Interpro were trained on the GO terms with at least one annotation. The results of the different implementations are shown in Table 4.

**Table 4.** The performance of different components or implementations of TransFew on the Test_all dataset in the three GO categories (BP, MF and CC). TransFew is the final version of the method in this work. Interpro, MSA, ESM, and TransFew+ stand for MLP (Interpro), MLP (MSA), MLP (ESM), and TransFew + MSA + Interpro, respectively.

| Methods | $F_{max} (\uparrow)$ | | | $WF_{max} (\uparrow)$ | | | $AUPR (\uparrow)$ | | | $S_{min} (\downarrow)$ | | |
| --- | --- | --- | --- | --- | --- | --- | --- | --- | --- | --- | --- | --- |
|  | BP | MF | CC | BP | MF | CC | BP | MF | CC | BP | MF | CC |
| Interpro | 0.368 | 0.632 | 0.667 | 0.330 | 0.604 | 0.547 | 0.281 | 0.596 | 0.729 | 29.809 | 7.219 | 7.494 |
| MSA | 0.372 | 0.628 | 0.671 | 0.332 | 0.600 | 0.549 | 0.285 | 0.592 | 0.677 | 29.616 | 7.504 | 7.635 |
| ESM | 0.515 | 0.690 | 0.735 | 0.478 | 0.662 | 0.638 | 0.471 | **0.654** | **0.788** | 22.875 | 6.367 | 6.551 |
| TransFew | **0.605** | **0.717** | **0.736** | **0.585** | **0.703** | **0.643** | **0.501** | 0.555 | 0.734 | **18.024** | **5.483** | **6.379** |
| TransFew+ | 0.351 | 0.679 | 0.716 | 0.296 | 0.659 | 0.617 | 0.235 | 0.497 | 0.629 | 30.323 | 6.265 | 6.964 |

Among the three methods of using only sequence representations to predict protein function, MLP (ESM) performs better than MLP (Interpro) and MLP (MSA) in terms of all the metrics for all three gene ontologies, indicating that the ESM2_t48 features are better than the MSA features and the Interpro features for generating sequence representations for protein function prediction.

TransFew that combines the sequence representation generated by ESM2_t48 and the label representation generated from the GO graph and the text description of GO terms performs better than MLP (ESM) without using the label representation in all but two situations, indicating that integrating the sequence representation and the label representation can generally improve protein function prediction.

It is a little surprising to observe that TransFew performs better than TransFew + MSA + Interpro that use the ESM2_t48 features with additional MSA and Interpro features to generate the sequence representation, suggesting that adding the MSA and Interpro features on top of the ESM2_t48 features does not improve the performance. The reason might be that adding the additional features does not increase relevant information much but the complexity of the model, leading to the deterioration of the generalization performance. Indeed, Supplementary Note 3 shows that TransFew + MSA + Interpro fits the training data better than TransFew but performs worse on the validation data than TransFew.

## 4. Conclusion and Future Work

In this work, we present a new approach (TransFew) combining the information in the input space (the protein sequence space) and the output space (the function label space) to improve protein function prediction, particularly the accuracy of predicting rare function terms (GO term labels with few annotations). In the input space, we use a large pretrained protein language model to generate features for a protein sequence, which are then mapped to the protein function space defined by both common and rare GO terms to create a function-relevant representation for the protein. Learning the unbaised representations of proteins in terms of both common and rare GO terms makes it possible to predict them on the equal footing. In the output space, we use a graph convolutional neural network-based auto-encoder to combine the textual definition of GO terms and the inheritance and composition relationships between GO terms in the GO graphs to generate the semantic representations for all the GO terms, capturing the similarity between GO terms to facilitate the transfer of annotation from common GO terms to rare ones.

The representations in the input space and the output space are integrated by a novel cross-attention mechanism to build the associations between the protein representation and the label representation, which are used to predict the final function terms for the protein. TransFew not only performs better than two highly sophisticated protein function prediction methods on newly released test proteins, but also can predict the rare GO terms more accurately, demonstrating the approach of representing and combining the data of GO terms and proteins is effective in predicting all kinds of GO terms.

Our experiment also demonstrates that the sequence features generated by a large protein language model (ESM2_t48) is sufficient to create a functional-relevant representation of protein sequences that is more useful for protein function prediction than the features generated from multiple sequence alignments (MSAs) and Interpro domain features. Even though this does not rule out the usefulness of MSAs and Interpro domain features, it does show that very large pretrained protein language models can effectively capture the evolutionary patterns in protein sequences relevant to protein function prediction.

Even though three modalities of data including protein sequences, textual description of GO terms, and hierarchical relationship between GO terms have been integrated by TransFew to predict protein function, other relevant modalities of protein data [43] such as protein structures, protein-protein interaction, hypothetical function annotations based on homology transfer, and the textual description of proteins have not be explored in this work. In the future, we plan to add all these modalities into our approach, harnessing the combined power of diverse features to further enhance the accuracy and robustness of protein function prediction. One promising avenue is to leverage multi-modal language models for protein function prediction, as demonstrated in computer vision [44, 45, 46] and a recent bi-modal protein language model [47]. By adopting such an approach, we can potentially learn a comprehensive representation of proteins encompassing protein sequence, structure, interaction, gene ontologies, and prior human knowledge to improve protein function prediction.

## 5. Acknowledgments

This work is supported in part by a grant from the National Science Foundation (NSF grant #: DBI2308699).

## References

1. UniProt Consortium. Uniprot: a worldwide hub of protein knowledge. Nucleic acids research, 47(D1):D506–D515, 2019.

2. Frimpong Boadu, Hongyuan Cao, and Jianlin Cheng. Combining protein sequences and structures with transformers and equivariant graph neural networks to predict protein function. bioRxiv, pages 2023–01, 2023.

3. The Gene Ontology Consortium. The Gene Ontology knowledgebase in 2023. Genetics, 224(1):iyad031, 03 2023.

4. Michael Ashburner, Catherine A Ball, Judith A Blake, David Botstein, Heather Butler, J Michael Cherry, Allan P Davis, Kara Dolinski, Selina S Dwight, Janan T Eppig, et al. Gene ontology: tool for the unification of biology. Nature genetics, 25(1):25–29, 2000.

5. Maxat Kulmanov and Robert Hoehndorf. DeepGOZero: improving protein function prediction from sequence and zero-shot learning based on ontology axioms. Bioinformatics, 38(Supplement 1):i238–i245, 06 2022.

6. Flood Sung, Yongxin Yang, Li Zhang, Tao Xiang, Philip HS Torr, and Timothy M Hospedales. Learning to compare: Relation network for few-shot learning. In Proceedings of the IEEE conference on computer vision and pattern recognition, pages 1199–1208, 2018.

7. Nabin Giri and Jianlin Cheng. De novo atomic protein structure modeling for cryo-em density maps using 3d transformer and hidden markov model. bioRxiv, 2024.

8. Kaveh Safavigerdini, Koundinya Nouduri, Ramakrishna Surya, Andrew Reinhard, Zach Quinlan, Filiz Bunyak, Matthew R Maschmann, and Kannappan Palaniappan. Predicting mechanical properties of carbon nanotube (cnt) images using multi-layer synthetic finite element model simulations. In 2023 IEEE International Conference on Image Processing (ICIP), pages 3264–3268. IEEE, 2023.

9. Ashwin Dhakal, Rajan Gyawali, Liguo Wang, and Jianlin Cheng. A large expert-curated cryo-em image dataset for machine learning protein particle picking. Scientific Data, 10(1):392, 2023.

10. Zheng Yuan and Doug Downey. Otyper: A neural architecture for open named entity typing. In Proceedings of the AAAI Conference on Artificial Intelligence, volume 32, 2018.

11. Tao Zhang, Congying Xia, Chun-Ta Lu, and Philip Yu. Mzet: Memory augmented zero-shot fine-grained named entity typing. arXiv preprint arXiv:2004.01267, 2020.

12. Farhad Pourpanah, Moloud Abdar, Yuxuan Luo, Xinlei Zhou, Ran Wang, Chee Peng Lim, Xi-Zhao Wang, and QM Jonathan Wu. A review of generalized zero-shot learning methods. IEEE transactions on pattern analysis and machine intelligence, 2022.

13. Yue Cao and Yang Shen. TALE: Transformer-based protein function Annotation with joint sequence–Label Embedding. Bioinformatics, 37(18):2825–2833, 03 2021.

14. Zeming Lin, Halil Akin, Roshan Rao, Brian Hie, Zhongkai Zhu, Wenting Lu, Nikita Smetanin, Allan dos Santos Costa, Maryam Fazel-Zarandi, Tom Sercu, Sal Candido, et al. Language models of protein sequences at the scale of evolution enable accurate structure prediction. bioRxiv, 2022.

15. Jinhyuk Lee, Wonjin Yoon, Sungdong Kim, Donghyeon Kim, Sunkyu Kim, Chan Ho So, and Jaewoo Kang. BioBERT: a pre-trained biomedical language representation model for biomedical text mining. Bioinformatics, 36(4):1234–1240, 09 2019.

16. Jacob Devlin, Ming-Wei Chang, Kenton Lee, and Kristina Toutanova. Bert: Pre-training of deep bidirectional transformers for language understanding. arXiv preprint arXiv:1810.04805, 2018.

17. Jinhyuk Lee, Wonjin Yoon, Sungdong Kim, Donghyeon Kim, Sunkyu Kim, Chan Ho So, and Jaewoo Kang. Biobert: a pre-trained biomedical language representation model for biomedical text mining. Bioinformatics, 36(4):1234–1240, 2020.

18. Thomas N Kipf and Max Welling. Variational graph auto-encoders. NIPS Workshop on Bayesian Deep Learning, 2016.

19. Roshan Rao, Jason Liu, Robert Verkuil, Joshua Meier, John F. Canny, Pieter Abbeel, Tom Sercu, and Alexander Rives. Msa transformer. bioRxiv, 2021.

20. Typhaine Paysan-Lafosse, Matthias Blum, Sara Chuguransky, Tiago Grego, Beatriz Lázaro Pinto, Gustavo A Salazar, Maxwell L Bileschi, Peer Bork, Alan Bridge, Lucy Colwell, et al. Interpro in 2022. Nucleic acids research, 51(D1):D418–D427, 2023.

21. Shaojun Wang, Ronghui You, Yunjia Liu, Yi Xiong, and Shanfeng Zhu. Netgo 3.0: Protein language model improves large-scale functional annotations. Genomics, Proteomics & Bioinformatics, 2023.

22. Shuwei Yao, Ronghui You, Shaojun Wang, Yi Xiong, Xiaodi Huang, and Shanfeng Zhu. Netgo 2.0: improving large-scale protein function prediction with massive sequence, text, domain, family and network information. Nucleic acids research, 49(W1):W469–W475, 2021.

23. Maxat Kulmanov and Robert Hoehndorf. Deepgozero: improving protein function prediction from sequence and zero-shot learning based on ontology axioms. Bioinformatics, 38(Supplement 1):i238–i245, 2022.

24. Adam Paszke, Sam Gross, Soumith Chintala, Gregory Chanan, Edward Yang, Zachary DeVito, Zeming Lin, Alban Desmaison, Luca Antiga, and Adam Lerer. Automatic differentiation in pytorch. 2017.

25. Adam Paszke, Sam Gross, Francisco Massa, Adam Lerer, James Bradbury, Gregory Chanan, Trevor Killeen, Zeming Lin, Natalia Gimelshein, Luca Antiga, et al. Pytorch: An imperative style, high-performance deep learning library. Advances in neural information processing systems, 32, 2019.

26. Thomas N. Kipf and Max Welling. Semi-supervised classification with graph convolutional networks. In International Conference on Learning Representations (ICLR), 2017.

27. Petar Veličković, Guillem Cucurull, Arantxa Casanova, Adriana Romero, Pietro Lio, and Yoshua Bengio. Graph attention networks. arXiv preprint arXiv:1710.10903, 2017.

28. Shaked Brody, Uri Alon, and Eran Yahav. How attentive are graph attention networks? arXiv preprint arXiv:2105.14491, 2021.

29. Yunsheng Shi, Zhengjie Huang, Shikun Feng, Hui Zhong, Wenjin Wang, and Yu Sun. Masked label prediction: Unified message passing model for semi-supervised classification. arXiv preprint arXiv:2009.03509, 2020.

30. Hanwen Xu and Sheng Wang. Protranslator: Zero-shot protein function prediction using textual description. In Research in Computational Molecular Biology: 26th Annual International Conference, RECOMB 2022, San Diego, CA, USA, May 22–25, 2022, Proceedings, page 279–294, Berlin, Heidelberg, 2022. Springer-Verlag.

31. Suzi A Aleksander, James Balhoff, Seth Carbon, J Michael Cherry, Harold J Drabkin, Dustin Ebert, Marc Feuermann, Pascale Gaudet, Nomi L Harris, et al. The gene ontology knowledgebase in 2023Genetics, 224(1):iyad031, 2023.

32. Predrag Radivojac, Wyatt T Clark, Tal Ronnen Oron, Alexandra M Schnoes, Tobias Wittkop, Artem Sokolov, Kiley Graim, Christopher Funk, Karin Verspoor, Asa Ben-Hur, et al. A large-scale evaluation of computational protein function prediction. Nature methods, 10(3):221– 227, 2013.

33. Martin Steinegger and Johannes Söding. Mmseqs2 enables sensitive protein sequence searching for the analysis of massive data sets. Nature biotechnology, 35(11):1026–1028, 2017.

34. Ronghui You, Shuwei Yao, Yi Xiong, Xiaodi Huang, Fengzhu Sun, Hiroshi Mamitsuka, and Shanfeng Zhu. NetGO: improving large-scale protein function prediction with massive network information. Nucleic Acids Research, 47(W1):W379–W387, 05 2019.

35. Wyatt T Clark and Predrag Radivojac. Information-theoretic evaluation of predicted ontological annotations. Bioinformatics, 29(13):i53–i61, 2013.

36. Naihui Zhou, Yuxiang Jiang, Timothy R Bergquist, Alexandra J Lee, Balint Z Kacsoh, Alex W Crocker, Kimberley A Lewis, George Georghiou, Huy N Nguyen, Md Nafiz Hamid, et al. The cafa challenge reports improved protein function prediction and new functional annotations for hundreds of genes through experimental screens. Genome biology, 20(1):1–23, 2019.

37. Yuxiang Jiang, Tal Ronnen Oron, Wyatt T Clark, Asma R Bankapur, Daniel D’ Andrea, Rosalba Lepore, Christopher S Funk, Indika Kahanda, Karin M Verspoor, Asa Ben-Hur, et al. An expanded evaluation of protein function prediction methods shows an improvement in accuracy. Genome biology, 17(1):1–19, 2016.

38. Yue Cao and Yang Shen. TALE: Transformer-based protein function Annotation with joint sequence–Label Embedding. Bioinformatics, 37(18):2825–2833, September 2021.

39. Benjamin Buchfink, Chao Xie, and Daniel H Huson. Fast and sensitive protein alignment using diamond. Nature methods, 12(1):59–60, 2015.

40. Damian Szklarczyk, Andrea Franceschini, Stefan Wyder, Kristoffer Forslund, Davide Heller, Jaime Huerta-Cepas, Milan Simonovic, Alexander Roth, Alberto Santos, Kalliopi P Tsafou, et al. String v10: protein–protein interaction networks, integrated over the tree of life. Nucleic acids research, 43(D1):D447–D452, 2015.

41. Quoc Le and Tomas Mikolov. Distributed representations of sentences and documents. In International conference on machine learning, pages 1188–1196. PMLR, 2014.

42. Alexander Rives, Joshua Meier, Tom Sercu, Siddharth Goyal, Zeming Lin, Jason Liu, Demi Guo, Myle Ott, C. Lawrence Zitnick, Jerry Ma, and Rob Fergus. Biological structure and function emerge from scaling unsupervised learning to 250 million protein sequences. PNAS, 2019.

43. Renzhi Cao and Jianlin Cheng. Integrated protein function prediction by mining function associations, sequences, and protein–protein and gene–gene interaction networks. Methods, 93:84–91, 2016.

44. Alec Radford, Jong Wook Kim, Chris Hallacy, Aditya Ramesh, Gabriel Goh, Sandhini Agarwal, Girish Sastry, Amanda Askell, Pamela Mishkin, Jack Clark, et al. Learning transferable visual models from natural language supervision. In International conference on machine learning, pages 8748–8763. PMLR, 2021.

45. Jean-Baptiste Alayrac, Jeff Donahue, Pauline Luc, Antoine Miech, Iain Barr, Yana Hasson, Karel Lenc, Arthur Mensch, Katherine Millican, Malcolm Reynolds, et al. Flamingo: a visual language model for few-shot learning. Advances in Neural Information Processing Systems, 35:23716–23736, 2022.

46. Julián N Acosta, Guido J Falcone, Pranav Rajpurkar, and Eric J Topol. Multimodal biomedical ai. Nature Medicine, 28(9):1773–1784, 2022.

47. Michael Heinzinger, Konstantin Weissenow, Joaquin Gomez Sanchez, Adrian Henkel, Martin Steinegger, and Burkhard Rost. Prostt5: Bilingual language model for protein sequence and structure. bioRxiv, pages 2023–07, 2023.

