## Supplementary Note 1 for "Improving protein function prediction by learning and integrating representations of protein sequences and function labels"

–

### Supplementary Materials

*Author: Frimpong Boadu, Jianlin Cheng*

### Supplementary Note 1: Generating MSA for a protein

We use HHblits to create a multiple-sequence alignment (MSA) for a protein sequence from the $UniRef30\_2022\_02$ database [4]. Subsequently, we filter out lowercase letters and insertion characters like ".", and "*" from the alignment. Employing a greedy algorithm[5], we choose 128 sequences that maximize the Hamming distance within the MSA. Finally, we employ the pre-trained ESM-MSA-1b [5] language model to generate representative embeddings for the protein using its MSA. The embeddings are used as input for the various MLP sub-models to generate representative embeddings for the protein.

### Supplementary Note 2: Generating Interpro domain features for a protein

We generate InterPro domain features from a protein sequence using InterProScan. Interpro features are represented as a binary matrix, where a 1 implies that a protein has an Interpro signature and 0 otherwise. In each of the three GO ontologies, we consider only signatures that appear in the training data, resulting in an input vector of dimension 24714, 25523 and 24846 for cellular component, molecular function and biological process respectively. The binary vector is then used as input for the various MLP sub-models to generate representative embeddings for the protein.

### Supplementary Note 3: Compare the training and validation process of TransFew and TransFew + InterPro + MSA

Figure 1 compares how TransFew and TransFew + InterPro + MSA behaved in the training and validation processes.

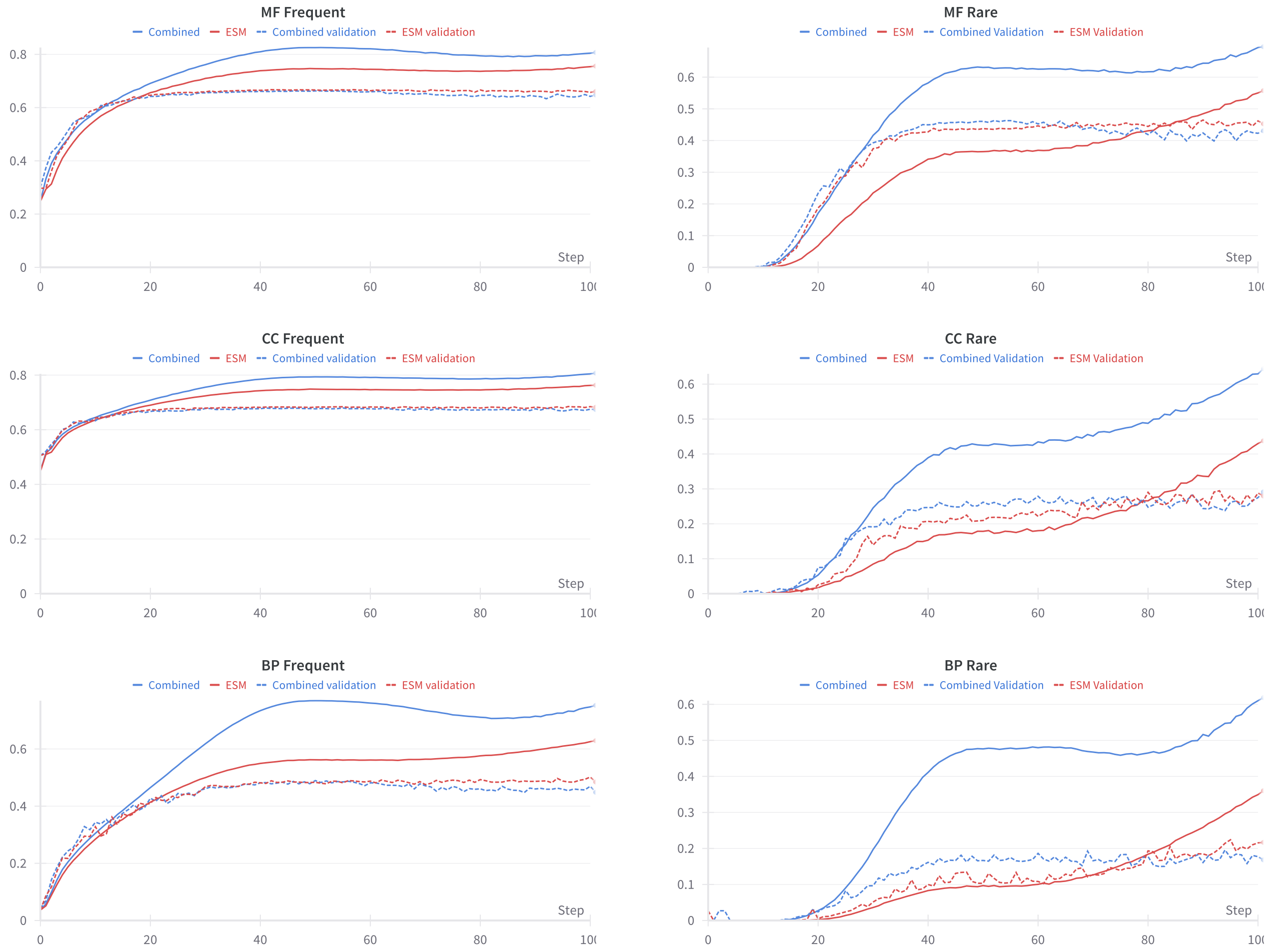


Figure 1: The training and validation curves for TransFew (in red) and TransFew + InterPro + MSA (called Combined). The sub-figures on the left are for the GO terms with annotation frequency greater than or equal to 30 (left) and the sub-figures on the right are for rare GO terms with annotation frequency with less than 30. Throughout the training process (solid lines), TransFew + InterPro + MSA consistently fits the training data better than TransFew, but on the validation data (dashed lines) Transfew performs better.

### Supplementary Note 4: Three label embedding methods

We explored three graph neural network-based auto-encoders to combine the features generated from the textual descriptions of GO terms using BioBERT and the ones from the hierarchical relationships between GO terms to create the representation of all GO terms, which are Graph Convolutional Network (GCN)-based auto-encoder[3], Graph Attention Networks (GAT)-based auto-encoder [1, 7], and Graph Transformer (TransformerConv)-based auto-encoder [6]. Their performance in the validation process is shown in Figure 2.

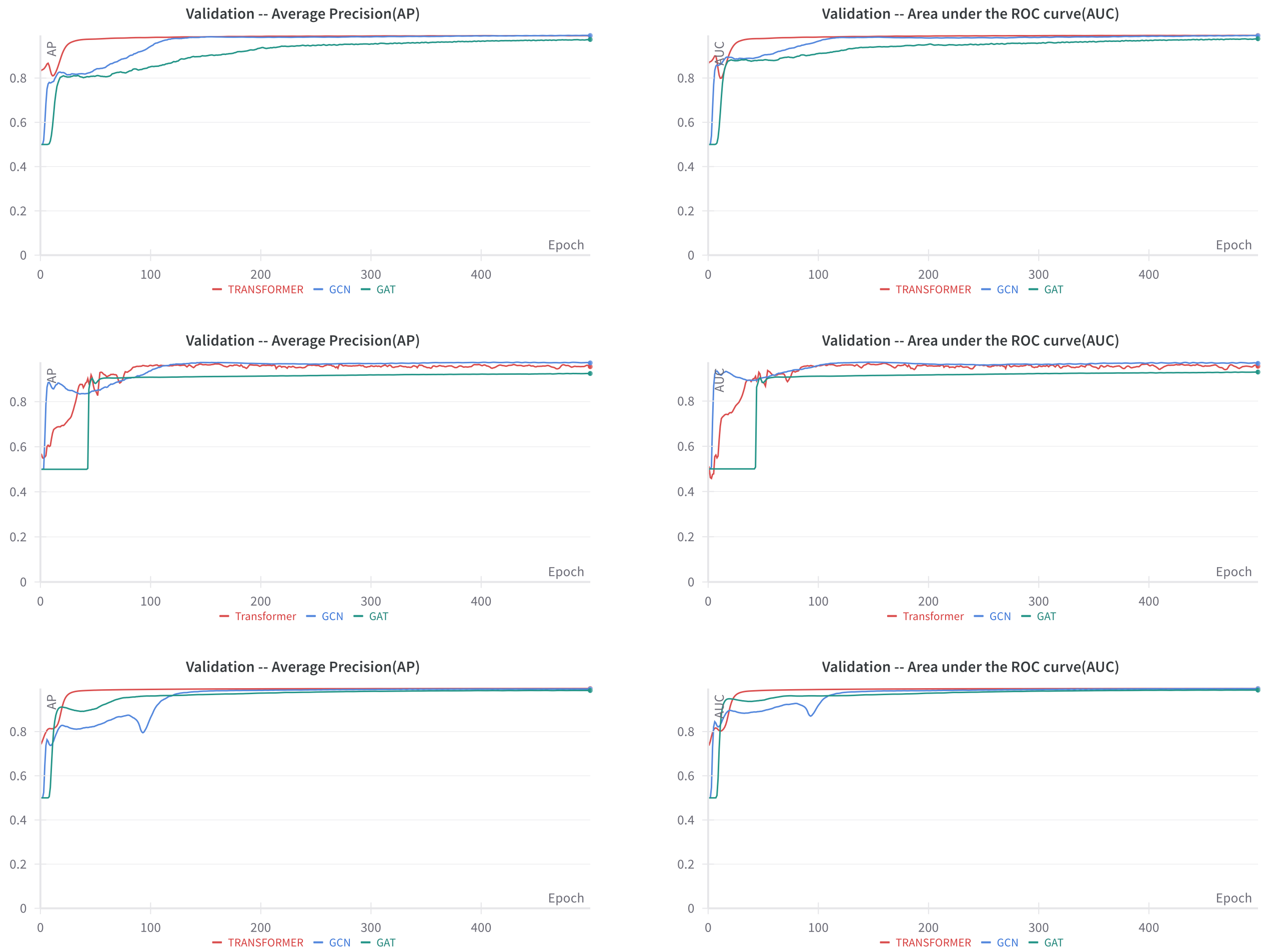


Figure 2: The Average Precision (AP) and Area under the ROC curve (AUC) of three graph neural network-based auto-encoders for three GO categories (molecular function (top), cellular component (middle), and biological process (bottom)). The three encoding architectures have similar performance..

### Supplementary Note 5: Evaluation Metrics

In this work, we use the three CAFA[2, 8] evaluation metrics: $F_{max}$, $S_{min}$, weighted $F_{max}$ and the area under the precision-recall curve (AUPR) to evaluate protein function predictions, which are defined as follows.

- **Precision**

$$\mathrm{pr}(\tau) = \frac{1}{m(\tau)} \sum_{i=1}^{m(\tau)} \frac{\sum_f \mathbb{I}(f \in P_i(\tau) \wedge f \in T_i)}{\sum_f \mathbb{I}(f \in P_i(\tau))}$$

- **Recall**

$$\mathrm{rc}(\tau) = \frac{1}{n_e} \sum_{i=1}^{n_e} \frac{\sum_f \mathbb{I}(f \in P_i(\tau) \wedge f \in T_i)}{\sum_f \mathbb{I}(f \in T_i)}$$

- **$\mathbf{F_1}$ Score**

$$F_{1(\tau)} = 2 \times \frac{\mathrm{pr}(\tau) \times \mathrm{rc}(\tau)}{\mathrm{pr}(\tau) + \mathrm{rc}(\tau)}$$

- **Maximum $\mathbf{F_1}$ Score**

$$F_{\max} = \max_{\tau} \left(F_1(\tau)\right)$$

  where $f$ is a term, $P_i(\tau)$ is the set of predictions, $T_i$ denotes the corresponding ground-truth, $i$ represents the protein sequence under consideration, and $\tau$ is the decision threshold. $m(\tau)$ is the number of proteins sequences with at least one predicted score greater than or equal to the decision threshold $\tau$, $\mathbb{I}(\cdot)$ is an indicator function, and $n_e$ is the number of proteins in the test set for a particular test study.

- **Information Content** ($ic$) of term $f$ is computed as:

$$\mathrm{IC}(f) = \log_2 \frac{1}{\mathrm{Pr}(f|P(f))}$$

- **Weighted precision**:

$$\mathrm{wpr}(\tau) = \frac{1}{m(\tau)} \sum_{i=1}^{m(\tau)} \frac{\sum_f ic(f) \cdot \mathbb{I}(f \in P_i(\tau) \wedge T_i(\tau))}{\sum_f ic(f) \cdot \mathbb{I}(f \in P_i(\tau))}$$

- **Weighted Recall**:

$$\text{wrc}(\tau) = \frac{1}{n_e}\sum_{i=1}^{n_e}\frac{\sum_f ic(f)\cdot \mathbb{I}(f \in P_i(\tau) \land T_i(\tau))}{\sum_f ic(f)\cdot \mathbb{I}(f \in T_i(\tau))}$$

  Here, $\Pr(f|P(f))$ represents the probability that term $f$ in the ontology is associated with a protein given that all of its parents are associated.

- **Remaining Uncertainty**

$$ru(\tau) = \frac{1}{ne}\sum_{i=1}^{ne}\sum_f ic(f)\cdot \mathbb{I}(f \notin P_i(\tau) \land f \in Ti)$$

- **Missing Information**

$$mi(\tau) = \frac{1}{ne}\sum_{i=1}^{ne}\sum_f ic(f)\cdot \mathbb{I}(f \in P_i(\tau) \land f \notin Ti)$$

- $S_{min}$

$$S_{min} = \min_{\tau}\sqrt{ru(\tau)^2 + mi(\tau)^2}, \tau$$

- **Area under precision recall curve (AUPR)**

$$\text{AUPR} = \int_0^1 \text{Precision}(R)\, dR$$

  where Precision(R) represents the precision at a given recall level (R).

### References


Brody, S., Alon, U., & Yahav, E. (2021). How attentive are graph attention networks? *arXiv preprint arXiv:2105.14491*.

Jiang, Y., Oron, T. R., Clark, W. T., Bankapur, A. R., D'Andrea, D., Lepore, R., … others (2016). An expanded evaluation of protein function prediction methods shows an improvement in accuracy. *Genome biology*, *17*(1), 1–19.

Kipf, T. N., & Welling, M. (2017). Semi-supervised classification with graph convolutional networks. In *International conference on learning representations (iclr).*

Mirdita, M., von den Driesch, L., Galiez, C., Martin, M. J., Söding, J., & Steinegger, M. (2016, 11). Uniclust databases of clustered and deeply annotated protein sequences and alignments. *Nucleic Acids Research*, *45*(D1), D170-D176. Retrieved from https://doi.org/10.1093/nar/gkw1081 doi: 10.1093/nar/gkw1081

Rao, R., Liu, J., Verkuil, R., Meier, J., Canny, J. F., Abbeel, P., … Rives, A. (2021). Msa transformer. *bioRxiv*. Retrieved from https://www.biorxiv.org/content/10.1101/2021.02.12.430858v1 doi: 10.1101/2021.02.12.430858

Shi, Y., Huang, Z., Feng, S., Zhong, H., Wang, W., & Sun, Y. (2020). Masked label prediction: Unified message passing model for semi-supervised classification. *arXiv preprint arXiv:2009.03509*.

Veličković, P., Cucurull, G., Casanova, A., Romero, A., Lio, P., & Bengio, Y. (2017). Graph attention networks. *arXiv preprint arXiv:1710.10903*.

Zhou, N., Jiang, Y., Bergquist, T. R., Lee, A. J., Kacsoh, B. Z., Crocker, A. W., … others (2019). The cafa challenge reports improved protein function prediction and new functional annotations for hundreds of genes through experimental screens. *Genome biology*, *20*(1), 1–23.
